# Linking the fundamental and realized niches in a hypersaline consumer-resource system

**DOI:** 10.64898/2026.06.26.734780

**Authors:** Lou Guyot, Stanislas Fereol, Roula Jabbour-Zahab, Luis-Miguel Chevin

## Abstract

A changing abiotic environment may affect the fitness and performance of an organism directly by challenging its environmental tolerance (related to its fundamental niche), or indirectly by modifying the abundance of other species it interacts with, and/or the strength of their interactions (related to its realized niche). The strength of an interaction may itself depend on how well each of the involved species performs across environments. However, there is a dearth of studies investigating how the intrinsic fitness of each interacting species covaries with the intensity of their interaction across an abiotic environmental gradient. We addressed this question in a hypersaline consumer-resource system: the microalga *Dunaliella* spp. grazed by the brine shrimp *Artemia franciscana*. We exposed four *Dunaliella* strains to a range of salinities, from just below to much above seawater, with or without brine shrimps. We tracked their population sizes over time and the survival of their grazers, to estimate basic parameters of a Lotka-Volterra model. We found that the intrinsic growth rate of algae, the survival rate of grazers, and the per-capita grazing rate, all varied with salinity and algal strain. Significant interactions between strain and salinity further revealed that these ecological responses to salinity are evolvable. Together with correlations between demographic parameters across salinity, this suggests that predation may influence the evolution of salinity tolerance curves, blurring the line between the fundamental and realized niches.

## Introduction

Understanding, predicting, and possibly controlling population responses to changing abiotic environments, such as rising temperature or drought, is becoming an urgent need in the current context of rapid anthropogenic global changes. There is a wealth of studies aiming to predict the impacts of such abiotic changes on species’ geographic distributions and extinction risk. This work ranges from purely statistical species distribution models (also described as environmental niche models; Guisan and Thuiller 2005) to more mechanistic approaches that explicitly account for demographic processes, sometimes combined with evolution and phenotypic plasticity (Crozier and Dwyer 2006; Kearney and Porter 2009). One limit that has been repeatedly put forward about these approaches is that they rarely take into account interactions with other species (Gilman et al. 2010; Urban et al. 2016; but see Pichler and Hartig 2021). There have been numerous calls in recent years for a tighter integration between population responses to the biotic and abiotic environments, to work out how a changing physical environment may affect species interactions, and reciprocally how species interactions matter for responses to abiotic environmental change (Gilman et al. 2010; Urban et al. 2016). This effort has been successfully initiated notably for competition across environmental gradients, with recent meta-analyses finding consistent support for a so-called ‘competitive exclusion–tolerance rule’, wherein a competitively dominant species excludes a subordinate species from benign environments, but the subordinate species then becomes better adapted to more stressful environments at the edge of the gradient (Martin and Ghalambor 2024, 2023).

Regarding trophic interactions, an important question at this interface is: To what extent are population responses to the abiotic environment in a focal species driven by direct effects of this environment on this species, versus indirectly mediated by the fact that this species tracks other species at lower trophic levels, which themselves respond to the changing abiotic environment? A textbook example illustrating this question is phenology, the seasonal timing of life history events. Phenology exhibits strong plastic and evolutionary responses to changing temperatures (Parmesan 2006; Radchuk et al. 2019), and also drives species interactions, with predators having to match the timing of their prey to maximize their fitness (and conversely prey having to avoid being synchronized with their predators) (Visser and Gienapp 2019; Samplonius et al. 2021). For instance, great tits use temperature as a cue to synchronize their reproduction with the peak in abundance of caterpillars that they use to feed their chicks (Visser and Gienapp 2019; Visser and Both 2005); caterpillars need to strike a balance between their temperature constraints (as ectotherms) and the availability of young tree leaves that they feed on (Asch et al. 2012; Phillimore et al. 2012); and trees need to precisely time their budburst so that their leaves and inflorescence are not destroyed by late frost in spring, while still completing their reproductive cycle before the next winter (Chuine and Beaubien 2001). Here, the responses to changing spring temperature are thus driven by direct responses to temperature at the lower trophic level (tree), but mostly by the match to the resource at the higher trophic level (birds). Beyond phenology, the abiotic environment may influence trophic interactions by affecting the degree of spatial overlap between prey and predators (Schweiger et al. 2008; Hunsicker et al. 2013), including depth in the water column for aquatic species, or by modifying (through evolution or plasticity) the traits that mediate their interaction strength (such as morphological or chemical defences; Agrawal 2001; Pančić and Kiørboe 2018; Lürling 2021; Harvell 1990). In other words, the intensity of consumer-resource (e.g. predator-prey) interactions along an environmental gradient is likely to depend on the extent of overlap between the fundamental niches of both interactors. However, this question has been little explored in a quantitative way, by measuring per-capita rates of population growth (or components thereof) across a gradient of an abiotic environment. Some studies have investigated how the details of the predatory interaction (e.g., functional response relating the number of consumed prey to their density, search or attack rate, handling time) are modified by temperature (Wang et al. 2022; Sentis et al. 2017, 2013, 2012; Veselý et al. 2019), but without relating these parameters to the intrinsic fitness of predators and prey, and usually focusing on binary environmental contrasts rather than more quantitative gradients (but see Sentis et al. 2012).

The relationship between the performances of predators and prey and their interaction strength across environments also bears on another important question: To what extent is evolution of the *fundamental* (rather than realized) niche influenced by ecological interactions? The fundamental niche of a species can be defined as the range of multi-dimensional environmental values over which its population growth is positive in the absence of ecological interactions (Hutchinson 1957). The realized niche in the field is then modified by interactions with other species, which either reduce (for antagonists such as predators, competitors or parasites) or expand it (for mutualists). A “slice” of the fundamental niche along one of its environmental axes (e.g. temperature, salinity) can be measured in the laboratory by assaying fitness components (survival and reproduction), or population growth rates (mostly for microbes), across values of the corresponding environmental variable, producing so-called environmental tolerance (or performance) curves (Cifuentes et al. 2001; Arnoldi et al. 2025; Lynch and Gabriel 1987). These tolerance curves are thought to be determined by (and evolve via) traits that mediate adaptation to the physical environment (Chevin et al. 2010; Lande 2014). However, gradients of an abiotic environment often coincide in the field with gradients of ecological interactions, with more benign environments being typically associated with more competitors and predators, while harsher environments may be associated with transitions from competition to mutualism (stress gradient hypothesis; Bertness and Callaway 1994). Because of such associations between abiotic and biotic environments, selective pressures are also likely to vary along abiotic gradients, for instance favoring avoidance of antagonists (defenses against grazers or predators) in more benign environments, but tolerance of abiotic stress in harsher environments. If this selection pattern has been sustained over the evolutionary history of a species, then the fundamental niche of this species may have evolved not only in response to the direct effect of the abiotic environment itself, but also to the type and intensity of ecological interactions encountered along this gradient. This can, in turn, influence current fitness responses to the abiotic environment, even when measured in the laboratory without interactors (environmental tolerance curves). A first step towards understanding whether and how much ecological interactions may mould evolution of the fundamental niche is to investigate: (i) how the strength of between-species interactions varies along abiotic environmental gradients; (ii) how this compares to environmental variation in the fitness of each species in the absence of interactions (tolerance curves, or fundamental niche); and (iii) whether genetic variance exists for these responses to the environment.

While the temperature responses of organisms, populations and communities have been explored in extensive detail (Angilletta 2009), including with respect to consumer-resource interactions (Wang et al. 2022; Sentis et al. 2017, 2013, 2012; Veselý et al. 2019), salinity has received comparatively much less attention. This is unfortunate, as salinity is changing at a rapid pace throughout the globe, with potentially dramatic ecological impacts (Lee et al. 2022). Ice melt is causing seawater to become fresher at high latitudes (with the Baltic Sea predicted to become close to a freshwater lake in less than a century), while salinity is rising at lower latitudes as a result of faster evaporation and reduced precipitation and riverine flow (Lee et al. 2022). The strong physiological impacts of salinity on water and ion balance (Lee et al. 2022) cascade up to influence community composition, especially in hypersaline environments such as salt lakes, coastal lagoons, and evaporation ponds, which are characterized by broad variation in salinity (Williams 1998; Oren 2009; Joint et al. 2002). This makes hypersaline systems ideal to study in detail how interaction strength varies relative to the fundamental niches of consumer and resource organisms. Zooplankton - phytoplankton interactions are particularly convenient to study in that regard, owing to their short generation times and small sizes. Some studies have investigated zooplankton grazing rates across salinity, including in hypersaline systems (Belovsky et al. 2024; Sura et al. 2017; Zadereev et al. 2022), but without using measurements that directly relate to explicit population dynamic definitions of the fundamental versus realized niche.

We here study the interaction between the microalga *Dunaliella* and the brine shrimp *Artemia* across salinity. *Dunaliella salina* is a halotolerant green microalga (Chlorophyceae), which can survive and reproduce across a broad range of salinities, from near freshwater to saturated brine (Oren 2005, 2014). It is the main primary producer in hypersaline environments, but this genus also includes the marine species *Dunaliella tertiolecta*, and the more hypersaline specialist *Dunaliella viridis* (Ben-Amotz 2009; Polle et al. 2020). *Artemia spp.* is a small crustacean of the class of branchiopods (like *Daphnia*), which can also tolerate a very broad range of salinity (albeit narrower than *Dunaliella*), from below to much above seawater, and is the main grazer in hypersaline environments (Browne 1991). *Dunaliella* and *Artemia* have a joint evolutionary history of interaction across a broad salinity range. Importantly, this interaction is largely exclusive in hypersaline environments, where *Dunaliella* is the only primary producer and *Artemia* the only grazer. We used this system to investigate how grazing intensity varies across salinity, and how this variation relates to the degree of overlap in the fundamental niches of interactors. Crucially, we strived to break down the correlation between abiotic and biotic gradients that typically exists in the field, where more negative interactions are generally found in more benign environments - here, more grazers at the lower salinity - by replenishing brine shrimp in environments where they died more. While this reduces the ecological realism of the experiment, it allows more firmly establishing causality, by distinguishing salinity effects on predation strength from its effects on the predator’s own environmental tolerance. We also considered within-and between-species variation in *Dunaliella* spp., to investigate the potential for evolution of a species’ fundamental niche as a result of varying interaction strength with its consumer along an environmental gradient. We interpret our experimental results in the light of classic trait-matching models of evolution of species interactions.

## Materials and Methods

### Strains and culture conditions

We used four strains of *Dunaliella* spp. Two strains of the halotolerant species *Dunaliella salina*, S1C and S13, were sampled in September 2020 from the Gruissan and La Palme salterns (respectively), in Southern France. Strain S1C was extracted from a salt crystal in the highest salinity pond, while strain S13 was isolated from a pond at lower salinity (17.75 on refractometer, equivalent to 3.42M NaCl). Strain S7V, also collected in Gruissan in September 2020 and isolated in April 2021, belongs to the sister species *Dunaliella viridis*, which is even more halophilic than *D. salina*, but unlike it does not turn orange through carotenogenesis at high light and salinity (Ben-Amotz 2009; Oren 2005; Borowitzka and Siva 2007). The last strain Ter belongs to the marine species *Dunaliella tertiolecta* (strain RCC6 from the Roscoff Culture Collection, provided by Giulia Ghedini in March 2023). All these strains were maintained in the laboratory at 24°C in artificial seawater complemented in salt to 2.4 M NaCl. Prior to the experiments, all four strains of *Dunaliella* were acclimated at all the assayed salinities. A few days before the experiment, the cultures were diluted to renew the medium and reduce the concentration, thus ensuring that the algae remained in exponential growth.

For the brine shrimp, we used the San Francisco Bay breed from the sexual species *Artemia franciscana*, obtained from dormant cysts collected in 1984 in Hayward, California, stored under dry conditions at 4°C. These cysts were hatched following the protocol described by Lievens et al. (2016). Briefly, they were rehydrated in deionized water for 2 to 3 hours, decapsulated in 2% sodium hypochlorite for a few minutes, and rinsed with water. Decapsulated cysts were then incubated in an aerated saline medium (artificial seawater adjusted at 0.4 M NaCl) at 28°C under constant light for a few days until emergence, before being acclimated under the conditions of the experiment detailed below.

### Grazing assay

The culture medium used for the experiments was custom-made artificial seawater, supplemented with NaCl to achieve the desired salinities (from 0.5 M to 3 M NaCl by steps of 0.5 M), and prepared following the recipes described by Zeballos et al. (2023, 2024) and Rescan et al. (2020). During the experiment, we cultured each of the 4 *Dunaliella* strains for 5 days at 24°C under light cycles of 12 hours with a light intensity of 200 μmol m^−2^ s^−1^, in 50 mL flasks (Cellstar, VWR 392-0016; Greiner Bio-One) containing 20 mL of medium, starting from a concentration of approximately 50,000 cells/mL. Our predation treatment included either no *Artemia* or 5 *Artemia* individuals, initially introduced as nauplius larvae. These larvae were first acclimated to the experimental conditions prior to the grazing assay. Newly hatched nauplii were placed at salinity 1 M for a few hours, then moved for 5 days to the different salinities used in the experiment (from 0.5 M to 3 M NaCl by steps of 0.5 M), with the S1C strain of *Dunaliella salina* as food source. Following this acclimation step, nauplius larvae were briefly rinsed in 50 mL medium of the same salinity, then placed with their corresponding *Dunaliella* strain and salinity for the grazing assay, with 4 replicates per condition, evenly split between two subsequent experimental batches. Throughout the experiment, the number of dead *Artemia* was recorded each day, and flasks with dead *Artemia* were replenished daily with individuals grown at salinity 1 M with S1C a food source, to maintain a constant number of 5 *Artemia* throughout. This procedure allowed decoupling the influence of salinity on *Artemia* survival from its effect on predation intensity per individual *Artemia*, in a way that would have been compromised if this predator had soon disappeared from environments that it does not tolerate well. Before entering the experiment, these replacement individuals were briefly rinsed and acclimated in 50 mL medium of the corresponding salinity.

### Algae concentration measurement

At the beginning of the experiment, then approximately every 24 hours over the next 4 days, each flask was agitated, and 200 µL of the medium was sampled to measure algae concentration using a Guava EasyCyte HT cytometer (Luminex Corporation, Texas, USA). Acquisition settings and gatings were set as described in Zeballos et al. (2023, 2024) and Rescan et al. (2020).

### Ecological and statistical models

We wish to estimate the parameters of a predator-prey (or consumer-resource) interaction based on Lotka & Volterra’s model (Kot 2001). This model consists of a pair of first-order differential equations for the dynamics of the abundances of prey (*N*) and predators (*P*),

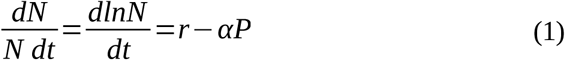

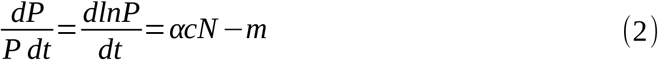

Equation (1) describes temporal changes in the population size (or density) of prey, where *r* is the intrinsic population growth rate of the prey (when at low density, without predators) and *α* is the *per-capita* predation rate. Equation (1) thus assumes that all density regulation in prey comes from predation, thereby neglecting the influence of within-species density dependence on population growth. This will be a good approximation as long as prey are kept at low density by their predators. Equation (2) describes temporal changes in the population size (or density) of predators, which additionally depends on the conversion rate *c* of prey into new predators, and the mortality rate *m* of predators. Here also, regulation of the density of predators is assumed to be entirely determined by the abundance of prey. Equation (2) also assumes a linear increase in predator population growth with the number of captured prey (type I functional response, as in the original Lotka-Volterra model). This is a good approximation as long as the abundance of prey does not saturate the handling time of predators, or their digestion capacity (Kot 2001). In short, this model describes the relationship between predator and prey populations, where the growth of prey is limited by predation, and the predator’s growth depends on the availability of prey (Kot 2001).

We are interested in characterising demographic responses to the environment (denoted with an index *ε*), which define the ecological niches of interactors (following Hutchinson’s 1957 definition), here with respect to salinity. As we are also comparing different algal strains, we additionally include an index *s* for strain. The fundamental niche of the prey is defined from variation in its intrinsic growth rate with the environment, which from equation (1) can be denoted as *r* (*ε, s*). For the predator, we considered that its fundamental niche is mostly determined by juvenile survival across environments when fed on the different strains, which we denote as 1−*m*(*ε, s*) (with *m* appearing in eq. 2). Finally, variation in the strength of predation (or grazing rate) among salinity and strains is captured by *α* (*ε, s*).

The population dynamic model in equations (1) and (2) was turned into a statistical model using generalised mixed models. For *Dunaliella,* we used the (rounded) concentration of algae (in numbers per mL) from day 1 to day 3 as the response variable, with a negative binomial error distribution and a log link function, consistent with Zeballos et al. (2023, 2024) and Rescan et al. (2020). The fixed effects included day, salinity, and salinity squared as continuous predictors, predation treatment (presence of 0 or 5 Artemia), and algae strain (S13, S1C, S7C, or Ter), with all interactions included (except for the interaction between salinity and salinity squared). The experimental batch was also included as a random effect. We estimated the parameters of this model using the glmmTMB package. In such a GLMM, an effect of day on cell concentration denotes a linear trend on the log scale, and therefore estimates an exponential growth (or decline) rate per unit day. This accurately described our experiment, where there was no evidence for density dependence in the early phase up to day 3 (Fig 1A). Interactions of day with other factors in turn quantified the effects of these factors on the exponential (i.e. per capita) growth rate. For instance, interactions of day with salinity and strain in the treatment without predators yield estimates of the dependence of the per-capita growth rate on salinity and strain, that is, *r* (*ε, s*) from eq. (1). Further including an interaction with predator number (effect of predation factor divided by 5, the number of predator) yields an estimate of *α* (*ε, s*), since it measures how each additional predator modifies the per-capita growth rate of prey. The interaction of the quadratic effect of salinity with day allows population growth rate to be maximised at an intermediate optimum salinity (if negative quadratic term; minimised otherwise). We additionally ran similar models (also using the glmmTMB package) separately for each condition of salinity, predation, and algae strain, which is identical to running a saturated model with one parameter per condition.

**Figure 1:**
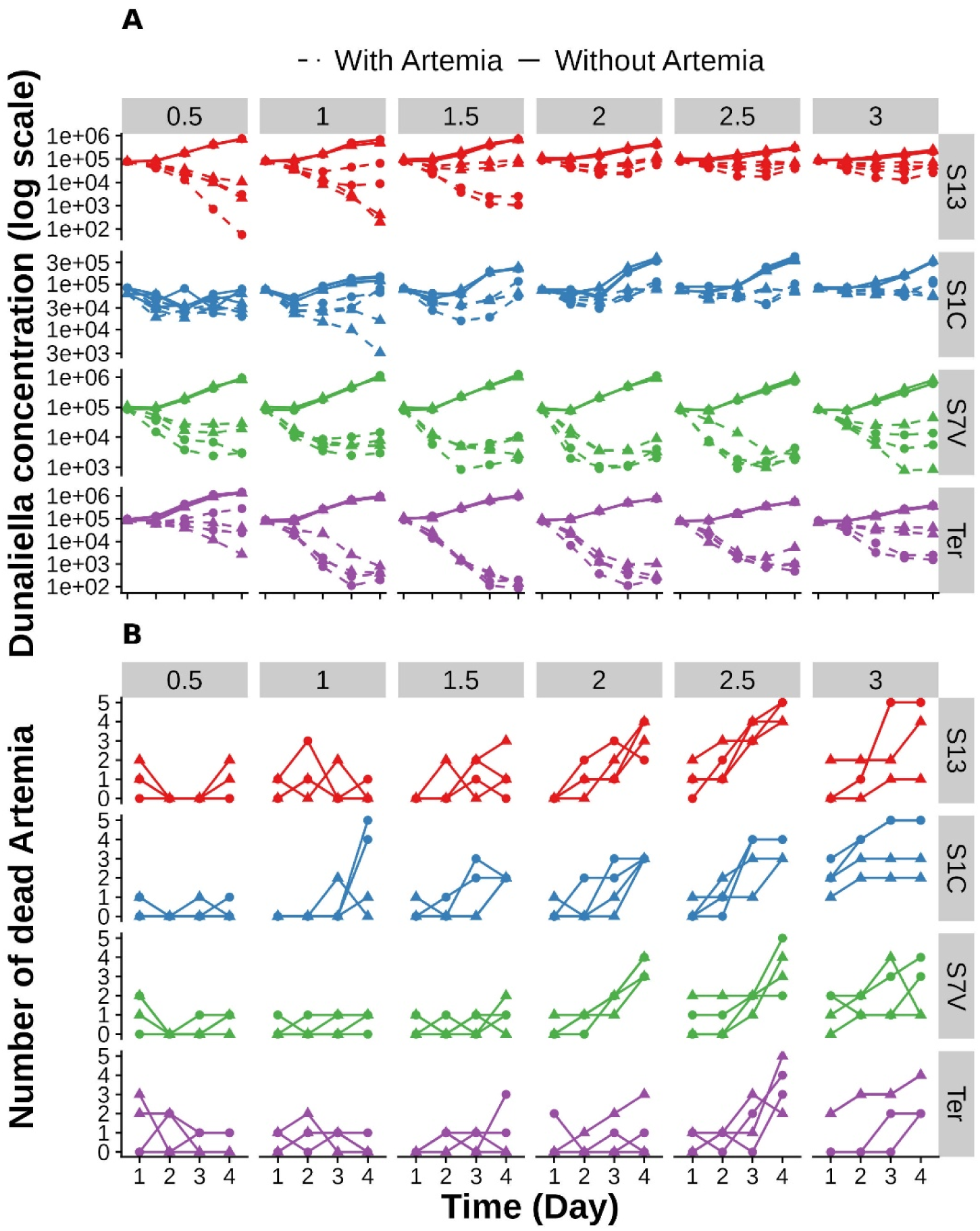
Demographic responses of *Artemia* and *Dunaliella* to variation in predation and salinity. A) Growth curves of *Dunaliella*, where log-transformed algal concentration measured by flow cytometry is plotted against measurement day, in the absence or presence of 5 *Artemia* nauplius larvae (continuous and dashed lines, respectively). B) Number of dead *Artemia* counted each day in each flask before their replacement. The legend above the graphs indicates the salinity concentration in M NaCl, and the legend on the right indicates the algal strains present in the flask. The dot shape corresponds to the experimental batch, with each batch composed of two replicate flasks. Colors refer to algal strain in both panels.

Brine shrimp survival probability in a flask was assessed using the number of nauplius larvae that survived until the end of the experiment (regardless of when they were added) and the total number that died during the experiment (i.e. cumulative death toll). We used logistic regression, that is, a generalised linear mixed-effects model, with the number of surviving and dead individuals as the response variables, a logit link function, and a binomial error distribution. Fixed effects included salinity (as continuous predictor), algal strain, and their interaction. Batch was treated as a random effect in the model. In such a setting, the effects of salinity and strains estimate respectively the effects of the environment and type of consumed algae on survival probability, while the interaction between salinity and strains quantifies how the type of algae consumed might affect salinity effects on survival (or reciprocally how salinity mediates the influence of the resource type on survival). The model was fitted with the lme4 package, using the maximum likelihood method with the Laplace approximation for random effects.

Predicted demographic parameters were obtained from the fitted models as linear combinations of fixed effects, according to the model structure. Standard errors were derived from the variance-covariance matrix of model coefficients using standard error propagation, and 95% confidence intervals were calculated as the estimate ± 1.96 SE (or after inverse-link transformation when appropriate). For *Dunaliella*, the per-capita predation rate α was derived as a contrast between predicted growth rates in predator-free and predator-present conditions divided by 5 (i.e. number of predators), assuming a linear reduction of prey growth rate with predator density as in eq. (1). The statistical significance of fixed effects was assessed using χ2-tests with the car package.

Relationships between the predicted demographic parameters were further examined using ordinary least-squares linear regressions across all strain × salinity combinations. The strength of the associations was quantified using the coefficient of determination (RZ), while statistical significance was assessed using the t-test associated with the regression slope, as provided by the fitted linear models.

Statistical analyses and graphic displays were performed using R software v 4.4.3 (R Core Team 2025), with the packages lme4 v 1.1.37 (Bates et al. 2015), glmmTMB v 1.1.13 (McGillycuddy et al. 2025), car v 3.1.3 (Fox and Weisberg 2019), and ggeffects v 2.3.1 (Lüdecke 2018). A statistical significance level of 5% was considered for all statistical tests. Data and codes used to generate figures and analyse data are available at: https://gitlab.com/Louguyot/env_tolerance_niche_realization_guyot_2026.

### Trait-dependent model of interaction and environmental tolerance

To help interpret these empirical results, we relied on classic models of phenotype-dependent interaction strength and environmental tolerance (Chevin and Chauhan 2025; Osmond et al. 2017; Guyot et al. 2026). These models were not used to derive new eco-evolutionary predictions (which would be beyond the scope of this paper, but see Osmond et al. 2017; Guyot et al. 2026); rather, they served to illustrate broad expected outcomes when phenotypic traits mediate both adaptation to the abiotic environment and interactions between species.

For adaptation to the abiotic environment, a typical assumption from such theory is that the changing environment causes movement of an optimum phenotype for selection (Lynch and Lande 1993; Bürger and Lynch 1995; Chevin et al. 2010; Kopp and Matuszewski 2014). We may thus write the prey’s growth rate in eq. (1) as 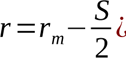, where *r_m_* is the maximal growth rate (for prey with well-adapted phenotype), *θ* is the corresponding optimum phenotype, and *S* is the strength of stabilising selection, such that the prey’s fitness decreases faster with increasing phenotypic deviation from the optimum when *S* is larger. The influence of the abiotic environment on this phenotype-to-fitness map is modeled by letting the optimum phenotype change linearly with the environment *ε*, that is, *θ* =*A* + *Bε*. The parameter *B* is the environmental sensitivity of selection (*sensu* Chevin et al 2010), which captures how fast the optimum phenotype for adaptation changes with the abiotic environment.

Regarding species interactions, a classic assumption from coevolutionary theory is that the *per-capita* strength of predation (*α* in eqs (1-2)) depends on the phenotypic match between the prey and the predator (Gavrilets 1997; Nuismer 2017; Gilman et al. 2012; Guimarães et al. 2017). We may thus write *α* =*Mexp* ¿, where *M* is the maximum predation strength (when the prey’s and predator’s traits match), *y* is the predator’s phenotype, and *γ* is a parameter that tunes how much predation strength changes in response to the phenotypic mismatch *x*− *y*. For larger *γ*, predation strength decreases more rapidly with increasing phenotypic mismatch.

Lastly, we let the traits of the predator and the prey exhibit some phenotypic plasticity in response to the abiotic environment. We assume a linear reaction norm for simplicity, such that the trait value of a prey exposed to environment *ε* is *x*=*a_x_* +*b_x_ ε* (and similarly *y*=*a_y_* +*b_y_ ε* for the predator). In this setting, the reaction norm slopes *b_x_* and *b _y_* quantify phenotypic plasticity for prey and predators, respectively, since they describe how fast their trait values change in response to the abiotic environment. The intercepts *a_x_* and *a _y_* determine the trait values in a reference environment.

## Results

To investigate how the fundamental niches of a prey and its predator relate to their interaction strength along an environmental gradient, we cultivated the microalga *Dunaliella* in the presence or absence of its main grazer *Artemia* under a broad range of salinities, from just below to much above seawater, and measured algae concentration over time. We also counted the living *Artemia* at each time point to assess their survival probability across conditions.

### Demographic responses of interactors

The population dynamics of *Dunaliella* without predators showed a strong repeatability between replicates (Fig 1A), but substantial variability between strains and, to a lesser extent, between salinities. More importantly, *Artemia*’s presence had a clear negative effect on the algae population growth. Indeed, algae populations tended to grow exponentially (following a short lag phase) in the absence of *Artemia*, while they declined drastically in the presence of *Artemia*, particularly from day 0 to 2, and showed more variability among replicates. Interestingly, a lag phase before initiation of population growth was observable in the early phase (between days 0-1) in the absence of *Artemia*, but was barely visible with *Artemia*. With *Artemia* nauplius larvae, a rebound in population growth was sometimes observed after day 2, which might be explained by an increased mortality of *Artemia* in the later part of the experiment. Consistent with this hypothesis, data on *Artemia* mortality confirmed that they died throughout the experiment, but with more individuals dying per day after day 2 and at high salinity (Fig 1B). Even though dead nauplii were replaced daily, the average number of living nauplii after day 2 was likely below five due to the high mortality rate. To limit the influence of these effects, in further analyses of population dynamics in *Dunaliella*, we worked on a subset of the data, keeping only days 1, 2 and 3, such that assuming exponential growth of *Dunaliella* (trend close to linear on the log scale in Fig 1A) and the presence of 5 *Artemia* remained reasonable.

### The fundamental niches of the prey and predator differ markedly

*Dunaliella* algae exhibited strain-specific salinity tolerance curves (Fig 2A). Their intrinsic growth rate varied across salinity (significant day x salinity interaction on algae concentration, Table S1) in a nonlinear manner (significant day x salinityZ interaction on algal concentration, Table S1). In addition, the intrinsic growth rate averaged across salinity significantly differed between *Dunaliella* strains (significant day x strain interaction on algae concentration, Table S1), as did the changes in this intrinsic growth rate with salinity (significant day x salinity x strain interaction on algae concentration, Table S1). The salinity tolerance breadth also differed between strains (significant day x salinityZ x strain interaction on algae concentration, Table S1). This suggests that there is evolutionary potential for salinity tolerance curves in *Dunaliella*. *Dunaliella viridis*, which is generally considered a high-salinity specialist, showed little variation in growth rate over salinity, but still had an optimal growth rate around 1.7 M NaCl (strain S7V, green in Fig 2A). The two strains from the halotolerant species *Dunaliella salina* (S13 and S1C, red and blue in Fig 2A) showed contrasted responses to salinity, with growth rate maximised respectively at low (0.5 M) and high (2 M) salinity. The strain from the marine species *Dunaliella tertiolecta* (Ter, purple in Fig 2A) grew better than *Dunaliella salina* at all salinities, but with an optimum at low salinity (0.5 M).

**Figure 2:**
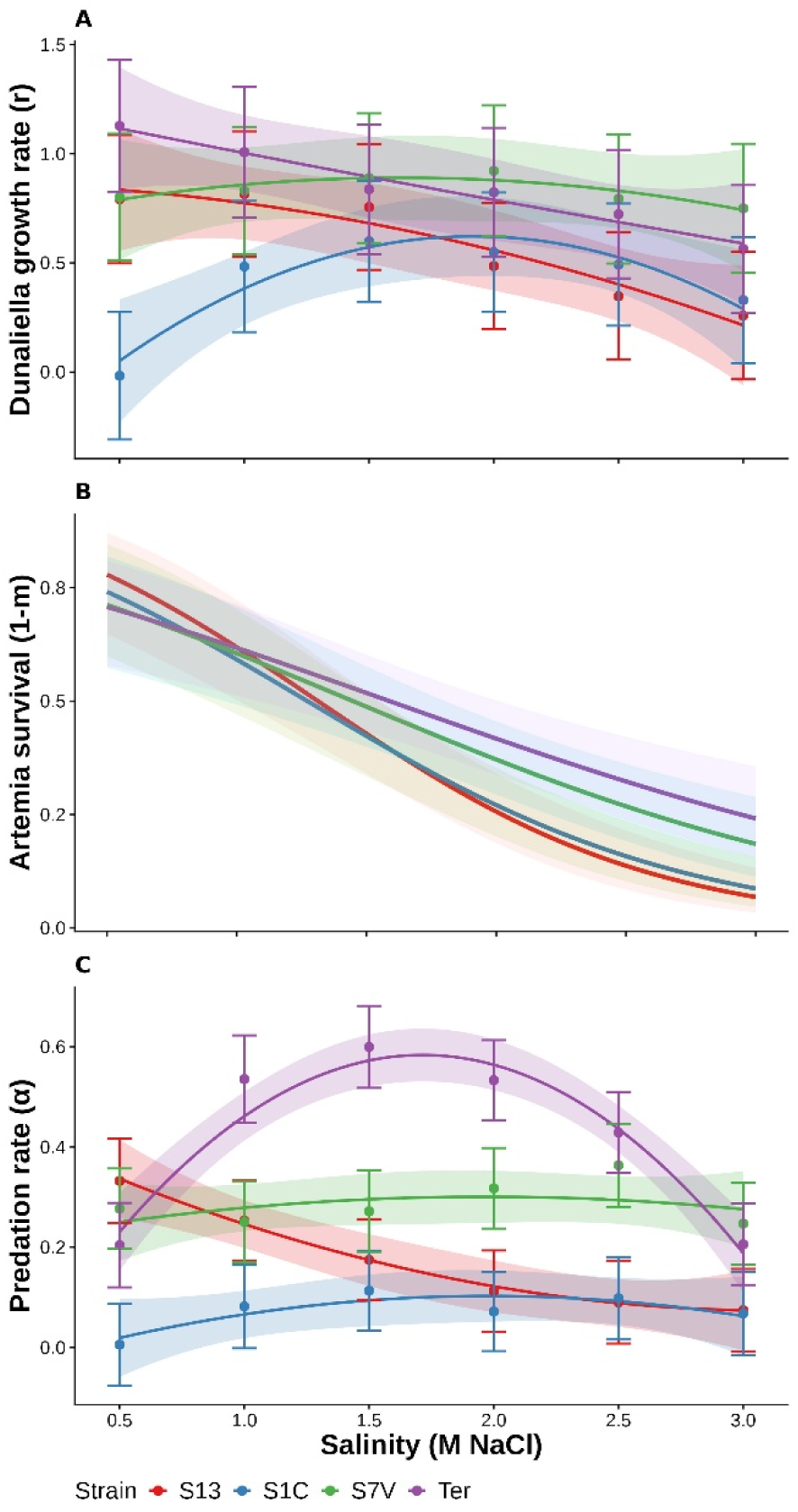
Fundamental niches and interaction strength of *Dunaliella and Artemia* across salinity. A) Salinity tolerance curves of *Dunaliella*, estimated from variation in the intrinsic growth rates of 4 *Dunaliella* strains across salinity. Dots and error bars represent estimates and standard errors from individual model fits for each strain and salinity, while curves and shaded error bands represent predictions and standard errors from a single model applied to the entire dataset, where salinity was treated as a continuous predictor, and strain and predation as categorical factors. B) Salinity tolerance curves of *Artemia*, estimating from the survival probability of nauplius larvae across salinity. Curves and shaded error bands represent model predictions and standard errors of a model where salinity is considered as a continuous predictor and strain as categorical factors. **C)** Variation in *per-capita* grazing rate across salinity, estimated from the difference between the growth rate of *Dunaliella* with and without *Artemia*, divided by 5 (i.e. the number of *Artemia*). Dots and error bars represent estimates and standard errors from individual model fits for each strain and salinity, while curves and shaded error band represent predictions and standard errors from a single model applied to the entire dataset, where salinity was treated as a continuous predictor, and strain and predation as categorical factors. Colors refer to algal strain in all panels.

In contrast, *Artemia franciscana* consistently survived better at lower salinities (Fig 2B). Regardless of the consumed algal strain consumed, increasing salinity substantially decreased the survival probability of *Artemia*, on average from 0.8 at 0.5 M NaCl to 0.1 at 3 M NaCl (significant effect of salinity on *Artemia* survival, Table S2), confirming that *Artemia* is more sensitive to salinity than its prey *Dunaliella*. However, the consumed *Dunaliella* strain influenced the steepness of this decrease in *Artemia* survival with salinity (significant salinity x strain interaction on *Artemia* survival, Table S2). Nevertheless, survival of *Artemia* averaged across salinity did not significantly differ between *Dunaliella* strains (no significant effect of strain on *Artemia* survival, Table S2). Hence, in this system, resource type (algae strain) had little overall impact on survival of the consumer (*Artemia*), but instead affected how this survival changed with salinity, or in other words, how well the consumer tolerated environmental stress.

### Interaction strength varies across salinity and strains

The per-capita predation rate varied across salinity (significant day x salinity x predation interaction on algal concentration, Table S1), in a nonlinear manner (significant day x salinityZ x predation interaction on algal concentration, Table S1). In addition, the mean predation rate (across salinities) depended on the strain (significant interaction day x strain x predation on algal concentration, Table S1; Fig 2C), with overall less predation on strains S1C and S13 than on S7V and Ter. Furthermore, how the predation rate changed with salinity depended on the algal strain (significant interactions day x strain x salinity x predation and day x strain x salinityZ x predation on algal concentration, Table S1). More specifically, strain Ter showed a highly non-linear response, with a maximum predation rate around 1.7 M (Fig 2C), while strains S1 and S7V had flatter responses, and predation rate decreased monotonically with salinity for S13. This suggests that there is evolutionary potential in *Dunaliella* for responses to *Artemia* predation across salinity.

A simple hypothesis for why predation intensity varies across salinity may be that predation efficiency is reduced in salinities where the predator *Artemia* has reduced performance. This can be investigated by using survival as a surrogate for overall performance, following Arnold (1983). However, the salinity where the predation rate was maximal appeared to differ from the salinity where *Artemia* survived best, especially for S7V, S1C and Ter (comparing Figs 2B and C). Accordingly, no pattern emerged when plotting the predicted predation rate against the predicted survival probability of *Artemia* across the strains and salinities investigated in our study (Fig 3A; non-significant correlation between predation rate and *Artemia* survival, Table S3A). Alternatively, predation intensity may have evolved to be maximal in environments where algal productivity is the highest. The salinity at which the *per capita* predation rate was maximised in Fig 2C largely matched the salinity at which *Dunaliella* grew best in Fig 2A (except to some extent for Ter). When considering all studied strains and salinities, the predicted predation rate indeed appeared higher in conditions associated with faster algal growth (Fig 3B; significant positive correlation between predation rate and algal growth rate, Table S3B). Note that, since *Dunaliella*’s growth rate without *Artemia* contributes to both axes of Fig 3B (see definition of *α* in the Materials and Methods), estimation error in this growth rate may contribute to the observed positive correlation. However, the mean variance of estimation errors for growth rates without *Artemia* was estimated at 0.012, corresponding to an estimated contribution of approximately 0.037 to the slope in Fig 3B. In addition, estimating *r* and *α* from different subsets of the data yielded similar results, indicating that the observed positive correlation is robust to potential effects of estimation error on the correlation (Supplementary Table 4). Combining both effects, the predation rate may depend on the niche overlap between partners, and thus be higher in conditions leading to both high productivity of the algal resource and high performance of the brine shrimp. This can be investigated by relating the predicted predation intensity to the product of the predicted *Artemia* survival by the predicted *Dunaliella* growth, as shown in Fig 3C. Again, across all strains and salinities investigated, predation intensity appeared to be higher where both *Artemia* survival and *Dunaliella* growth were high (Fig 3C; significant positive correlation between predation rate and the product between brine shrimp survival and algae growth rate, Table S3C).

**Figure 3:**
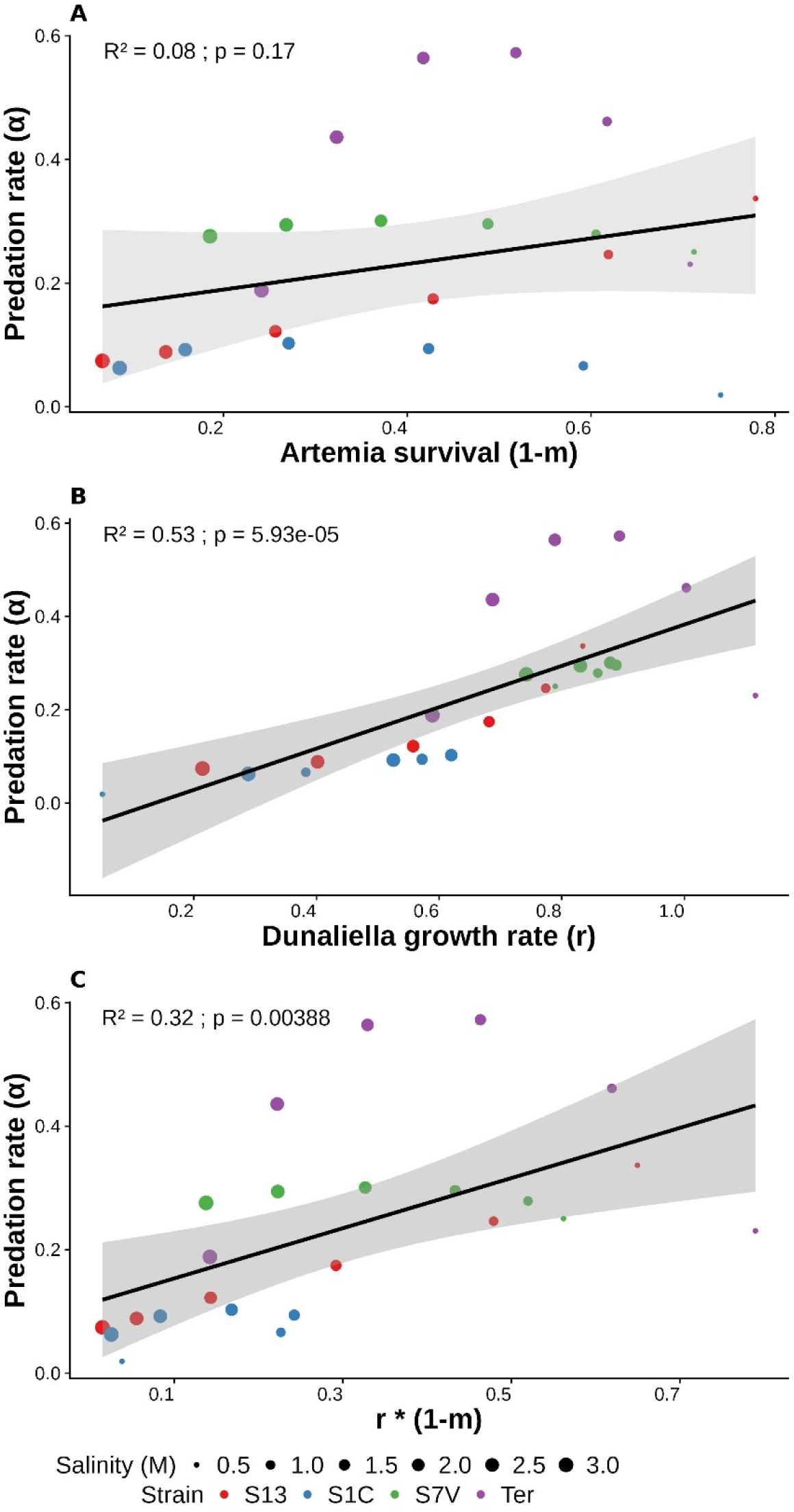
Relationship between the fundamental niches and interaction strength across salinity. Model predictions for the grazing rate are plotted against the predictions for *Artemia* survival (A), *Dunaliella* intrinsic growth rate (B), or their product (C), across each strain at salinity investigated in the experiment. Dot colors refer to algal strains, and dot size corresponds to salinity, ranging from 0.5 to 3 M of NaCl. The coefficients of determination (RZ) and p-values associated with the regression slope (p) are indicated on the top left.

### Phenotype-matching model suggests plasticity of interaction traits

These experimental results can be interpreted in the light of theory where both adaptation to the abiotic environment and species interactions are mediated by the same (or correlated) phenotypic traits (Chevin and Chauhan 2025; Guyot et al. 2026; Osmond et al. 2017). Broad insights can be gained by combining three key ingredients from classic models of phenotypic evolution: (i) a moving optimum phenotype, whereby the trait value that maximizes intrinsic fitness (for prey and/or predators) changes with the environment; (ii) trait-matching interaction, such that the predation rate is highest when predators and prey have phenotypes that match; and (iii) phenotypic plasticity, allowing the traits of predators and prey to directly respond to the environment. Under these classic assumptions, the plasticity of the prey determines its tolerance breadth, that is, how fast its intrinsic growth rate decreases when moving away from the optimal environment (Fig 4A, analogous to empirical Fig 2A-B), as previously pointed out by Chevin et al. (2010). Interestingly, these models also predict that the breadth of the curve relating interaction strength to the environment should depend on the difference in plasticity between the prey and the predator (Figure 4B, analogous to Fig 2C). In particular, when plasticity is the same in the prey and the predator (*b_x_*=*b_y_*), including when neither is plastic (*b_x_*=*b_y_*=0), a matching-trait model of interaction predicts that predation intensity should not vary with the environment (green curve in Fig 4B), because the phenotypic mismatch between prey and predators remains constant across environments. When considering both aspects of the niche jointly, plasticity in the prey and predators can profoundly impact the relationship between *r* and *α* across the abiotic environment (Fig 4C, analogous to Fig 3B). While Figure 4 is only illustrative and is far from an exhaustive theoretical exploration, it serves to illustrate that phenotypic plasticity not only influences the fundamental niche of a species (as captured by environmental tolerance curves, Chevin et al. 2010), but also how this fundamental niche differs from the realized niche (as captured by environmental changes in the strength of ecological interaction), and how these two aspects are related across environments.

**Figure 4:**
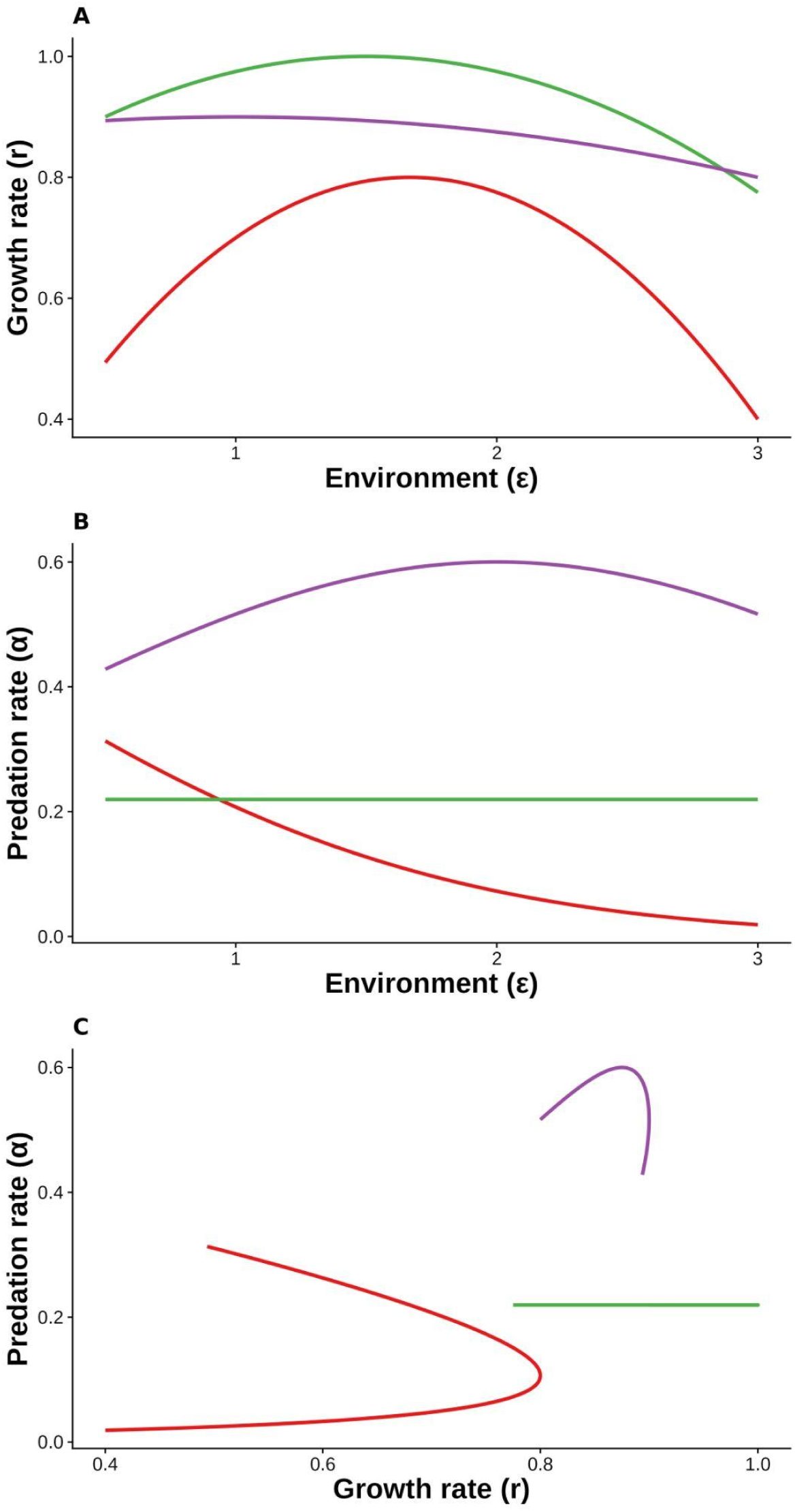
Predictions from a trait-matching model of interaction. The growth rate *r* of prey (A) and the predation intensity *α* (B) are represented along the environment, for a model where the environment determines the optimum phenotype for a trait in the prey, and the match between this trait and a corresponding trait in the predator determines the predation intensity. The relationship between *r* and *α* across environments is also represented in C. The involved traits are phenotypically plastic, with plasticity in the prey being lower (red), equal (green), or higher (purple) than plasticity in the predator.

## Discussion

### Relating fundamental niches to interaction strength

We quantified the fundamental niches of the microalga *Dunaliella* and its grazer, the brine shrimp *Artemia,* to compare them with the strength of their interaction across salinity. A first striking observation was that *Dunaliella* and *Artemia* had markedly different responses to salinity: while the brine shrimp systematically survived better at lower salinities (Fig 2B), some *Dunaliella* strains (S1C and S7V) grew best at the intermediate salinities we assayed (1.5 to 2 M NaCl; Fig 2A), which represent hypersaline environments much above seawater (∼0.6 M NaCl). In fact, the intrinsic growth rate of *Dunaliella* without predators changed little across the broad salinity range that we investigated. While *D. salina* (S1C and S13) and *D. viridis* (S7V) are halotolerant species that are expected to show little sensitivity to salinity, it is perhaps more surprising that the marine species *D. tertiolecta* (Ter) still grew reasonably well at the highest salinity in our assay (3 M = 175 g/L NaCl). However, this is consistent with the fact that the marine status of *D. tertiolecta* is a derived state, in a group that is otherwise restricted to hypersaline environments (Henley et al. 2018; Assunção et al. 2012, 2013). It would be worthwhile investigating the outcome of competition between *D. tertiolecta* and the hypersaline species *D. salina* and *D. viridis* in marine versus hypersaline environments, to test whether they conform to the ‘competitive exclusion–tolerance rule’ that was recently put forward by Martin and Ghalambor (2024, 2023).

Another important observation was that the *Dunaliella* strains differed in how they impacted the survival (Fig 2B) and grazing rate (Fig 2C) of *Artemia*, and how these impacts varied across salinity. The differences were more marked for the grazing rate, which was overall highest for Ter and lowest for S1C across salinities, and went from nearly flat (S1C and S7V) to declining (S13), to curved with a maximum at an intermediate salinity (Ter). As predation rate is an interaction trait, neither its average value nor its variation across strains and salinity can be firmly attributed to traits of the predator or of the prey without further investigation. On the prey side, *Dunaliella* strains may exhibit different constitutive and inducible defense mechanisms against grazing, from swimming at different depth to forming aggregates or producing toxic compounds (Lürling 2021; Pančić and Kiørboe 2018), and these mechanisms may vary with salinity. On the predator side, *Artemia* may limit its activity in more stressful environments (i.e., at higher salinities), thus limiting its food intake, which is directly related to movement in such filter feeders (Belovsky et al. 2024; Sura et al. 2017; Zadereev et al. 2022). Differences in grazing rates among *Dunaliella* strains may also be partly attributable to *Artemia*, which was shown to be able to graze selectively (Belovsky et al. 2024), and thus to exhibit preference in its filtering behavior. Nutritive value might also play a role, as *D. salina* is the eucaryotic organism that can produce the most beta-carotene per dry weight (Oren 2005; Ben-Amotz 2009), while *D. viridis* and *D. tertiolecta* do not produce this very nutritious lipid. Accordingly, the most orange *D. salina* strain S1C was less consumed than others and thus barely declined in the presence of *Artemia* (dashed blue lines in Fig 1A), despite growing more slowly when alone (continuous blue lines Fig 1A, and blue line in Fig 2A). This pattern is consistent with the hypothesis that S1C satiates *Artemia* faster, thus limiting the grazing rate on this strain.

While investigating these proximate mechanisms of the interaction is beyond reach in this study, we instead focused on a simpler hypothesis that could be addressed with our experimental design: that the variation in interaction strength (grazing rate) among strains and salinities was partly explained by the joint performances of both partners. Using fitness components as proxies for performance (following Arnold 1983), we indeed found that the predation rate was higher in conditions where the *Dunaliella* strains grew faster (Fig 3B), and (to a lesser extent and non-significantly) where *Artemia* survived better (Fig 3A). When considering both terms together, grazing rate was higher in conditions where both the prey and predator performed better (Fig 3C). This may either reflect some broad physiological constraints that jointly affect the performance of both partners and their interaction strength, or instead result from specific predator-prey coevolution for traits that trade off with adaptation to the abiotic environment (Guyot et al. 2026).

In an attempt to interpret these findings in the light of broader concepts in predator-prey interactions and coevolution across environments, we turned to the classic theoretical frameworks of trait-matching interactions and moving optimum models. While a thorough investigation was beyond reach in this experimental study, a key finding was that, under trait-matching interaction, variation in predation intensity across abiotic environments (here salinity) is only expected to occur if the predator and prey differ in their plastic responses to this environment (Fig 4B). As phenotypic plasticity is also expected to be a key determinant of environmental tolerance breadth (Chevin et al. 2010; Lande 2014), it is likely to play a key role in the links between the fundamental and realized niches, and how they jointly evolve in variable environments.

### Limits and perspectives

While we strived to estimate the fitness components of prey and predator as accurately as possible, these estimates were inherently constrained by our experimental design. Regarding the population growth rate of *Dunaliella*, we had to restrict our analyses to the time window between days 1 and 3, for consistency between the treatments with and without predation. Indeed, in conditions without predation, a lag phase was generally observed prior to that (from day 0 to 1), manifested as a flat part of the growth curve (dashed lines in Fig 1A). And in conditions with predation, later days (from 3 to 4) generally showed a rebound in population growth (dashed lines in Fig 1A), probably due to excess mortality of *Artemia* (Fig 1B). Days 1 to 3 were the best compromise to yield accurate estimates of the growth rate of *Dunaliella* with versus without predation, while reducing the impact of these two limits, but they still included part of the lag or rebound phases under some conditions.

Regarding *Artemia* survival, our experiment was not optimally designed for its accurate estimation, which would have required growing individuals in separate vials to limit competitive effects, and tracking the fate of these individuals to estimate parameters of a survival model (e.g. hazard rates and Kaplan Meier curves), as performed with *Artemia* across salinity by Nougué et al. (2016). Nevertheless, our procedure of replenishing dead *Artemia* individuals was efficient in allowing us to maintain a relatively stable grazing pressure throughout the experiment, while also providing a large enough number of *Artemia* individuals from which to estimate mortality rates. A caveat is however that variation in condition and age of brine shrimps probably contributed to: (i) the large variation observed between replicate *Dunaliella* populations in conditions with predation (spread of dashed lines in Fig 1A); and (ii) the rebound found in later days in days under these conditions, which coincides with increased daily mortality of *Artemia* (Fig 1B). Another obvious limitation of this design is that replenishing consumers in environments where they would normally die lacks ecological realism. However this procedure was not designed to mimic natural conditions, but rather to allow accurate estimation of environmental effects on grazing rates, while disentangling them from the environmental tolerance of the grazer. Nevertheless, the parameter values estimated through this protocol could still lead to realistic ecological projections if inserted in the consumer-resource system in equations (1-2); for instance, they would correctly predict that predators have weaker influence on prey in environments that they tolerate less well (i.e. at the high salinities where they die more).

Other implicit assumptions in our analysis had to do with the ecological underpinnings of the interaction. First, we assumed that *Artemia* exerted a constant *per-capita* grazing pressure throughout the experiment, thus ignoring that surviving nauplius larvae grew larger during the experiment. However, a 4-day experiment was likely short enough relative to the life cycle of this species to neglect putative effects of developmental stage on grazing (Browne 2018; Reeve 1963). On the other hand, the increased mortality of *Artemia* in later days and at high salinities (Fig 1B) meant that fewer individuals were grazing under these conditions on average (despite daily replacement of dead individuals), thus reducing the net predation rate and leading to a rebound in *Dunaliella* concentration after day 2-3 (dashed lines in Fig 1A). We also assumed that *Dunaliella* was sufficiently abundant that the grazing rate was maximum and constant during the time interval over which we estimated it. This is compatible with the observation that *Dunaliella* was not depleted by the end of the experiment, despite reaching very low concentrations in some conditions (Fig 1A). We also implicitly assumed that *Artemia* was not fully satiated such that it stopped grazing after some time. It would be interesting to pursue such an experiment for longer, allowing *Dunaliella* to reach a stationary population size (carrying capacity) in conditions without predation, and a much lower stable population size at a balance between grazing and growth in conditions with predation, to estimate the influence of predation on equilibrium population dynamics, which may be more relevant to natural conditions that are relatively stable.

Another critical assumption underlying our inference of predation rates was that the basic demographic parameters of algae, especially their intrinsic rates of increase, were identical with and without predation, consistent with most predation prey models (Kot 2001). This assumption may however not be consistent with the fact that a plateau was observed in the first day (lag phase) without *Artemia*, while no slope break was visible with *Artemia*. The assumption of a constant intrinsic rate of increase with and without predation may be violated in particular if plastic inducible defenses against predation exist in *Dunaliella*, and exhibit plasticity costs (DeWitt 1998; Dewitt et al. 1998). Studies that investigated plastic responses to predation in phytoplankton species by exposing them to predator cues have demonstrated that these responses are indeed common, and can be costly (Lürling 2021; Pančić and Kiørboe 2018). Note that some inducible defenses may take a few days to mount, as observed for the plasticity of other traits in this system (Leung et al. 2020; Cheloni and Slaveykova 2021), which may partly explain the increased death rate of *Artemia* towards the end of the experiment. Investigating such inducible responses to predation across salinity would help disentangle the mechanisms behind the co-variation in the fundamental and realized niche in this system.

### Conclusion and perspectives

Leveraging on a planktonic consumer-resource system, and using the ecological formalism of the Lotka-Volterra predator-prey model, we showed that, despite marked differences between the fundamental niches of predators and prey, the predation rate was highest where these niches overlapped. Combined with genetic variation in the prey for these joint responses to salinity and predation, this suggests that interactions with predators across salinity may impact the evolution not only of the realized niche of prey, but also of their fundamental niche. This hypothesis could be investigated using experimental evolution, where predation intensity is manipulated together with salinity. Identifying the traits involved in the interaction, some of which are well described for microalgae (Lürling 2021; Pančić and Kiørboe 2018), would help relate these experimental findings to the predictions from predator-prey coevolutionary theory in a changing abiotic environment (Guyot et al. 2026; Osmond et al. 2017; Jones 2008). Ultimately, the combination of laboratory experiment (including experimental evolution), theoretical predictions, and analyses of natural populations, will be crucial in order to understand and predict the responses of species to joint changes in their biotic and abiotic environments.

## Acknowledgements

We thank an anonymous reviewer for useful feedback.

## Author contributions

RZ provided the Artemia cysts and trained SF on cyst hatching and raising of nauplii. LMC, SF and LG designed experiments. LG and SF performed experiments. LMC supervised the study. LG analysed the data. LG produced the figures. LMC and LG wrote the manuscript.

## Supplementary tables

**Supplementary Table 1:** Analysis of deviance for rounded *Dunaliella* concentration (per mL). A generalised linear mixed model (GLMM) was used, with a negative binomial distribution and log link function. Fixed effects include day (from 1 to 3, treated as continuous), salinity (linear and quadratic terms i.e., salinityZ), predation treatment (0 or 5 *Artemia* nauplii), and strain (S13, S1C, S7C, or Ter), with all interactions among predictors, including linear and quadratic salinity terms. Note that all interactions with day denote an effect on the *per-capita* growth rate of the population. Batch was included as a random intercept. Effects were tested using Type II Wald chi-square tests. Columns report the chi-square statistic (Chisq), degrees of freedom (Df), and associated p-values (Pr(>Chisq)). Significance levels are indicated as follows: *** p < 0.001, ** p < 0.01, * p < 0.05,. p < 0.1.

| **Effect** | **Chisq** | **Df** | **Pr(>Chisq)** |
|---|---|---|---|
| day | 5.8044 | 1 | 0.01599 * |
| salinity | 113.3760 | 1 | < 2.2e-16 *** |
| salinity<sup>2</sup> | 108.3480 | 1 | < 2.2e-16 *** |
| predation | 3521.7581 | 1 | < 2.2e-16 *** |
| strain | 183.4057 | 3 | < 2.2e-16 *** |
| day:salinity | 6.2364 | 1 | 0.01251 * |
| day:salinity<sup>2</sup> | 4.6657 | 1 | 0.03077 * |
| day:predation | 697.4383 | 1 | < 2.2e-16 *** |
| salinity:predation | 177.4055 | 1 | < 2.2e-16 *** |
| salinity<sup>2</sup>:predation | 196.8932 | 1 | < 2.2e-16 *** |
| day:strain | 57.9384 | 3 | 1.620e-12 *** |
| salinity:strain | 289.7362 | 3 | < 2.2e-16 *** |
| salinity<sup>2</sup>:strain | 212.6518 | 3 | < 2.2e-16 *** |
| predation:strain | 878.4400 | 3 | < 2.2e-16 *** |
| day:salinity:predation | 18.1096 | 1 | 2.085e-05 *** |
| day:salinity<sup>2</sup>:predation | 22.8117 | 1 | 1.787e-06 *** |
| day:salinity:strain | 101.5415 | 3 | < 2.2e-16 *** |
| day:salinity<sup>2</sup>:strain | 90.1301 | 3 | < 2.2e-16 *** |
| day:predation:strain | 201.3449 | 3 | < 2.2e-16 *** |
| salinity:predation:strain | 191.5327 | 3 | < 2.2e-16 *** |
| salinity<sup>2</sup>:predation:strain | 173.4635 | 3 | < 2.2e-16 *** |
| day:salinity:predation:strain | 58.7485 | 3 | 1.088e-12 *** |
| day:salinity<sup>2</sup>:predation:strain | 54.8757 | 3 | 7.298e-12 *** |

**Supplementary Table 2:** Analysis of deviance for Artemia survival. A generalised linear mixed model was used with a binomial distribution and logit link function. Fixed effects include salinity, strain, and their interaction. Batch was included as a random intercept. Effects were tested using Type II Wald chi-square tests. Columns report the chi-square statistic (Chisq), degrees of freedom (Df), and associated p-values (Pr(>Chisq)). Significance levels are indicated as follows: *** p < 0.001, ** p < 0.01, * p < 0.05,. p < 0.1.

| **Effect** | **Chisq** | **Df** | **Pr(>Chisq)** |
|---|---|---|---|
| salinity | 114.2204 | 1 | < 2e-16 *** |
| strain | 6.9838 | 3 | 0.07242 . |
| salinity:strain | 7.8622 | 3 | 0.04895 * |

**Supplementary Table 3:** Linear regression models for the relationships between predicted demographic parameters across strains and salinities. Models evaluate the relationship between the predicted predation rate and A) the predicted *Artemia* survival probability, B) the predicted *Dunaliella* growth rate, and C) the predicted product *Artemia* survival × *Dunaliella* growth rate. For each model, regression coefficients (β), standard errors (SE), t-values, and associated p-values are reported. Model fit is reported using the coefficient of determination (RZ). Significance levels are indicated as follows: *** p < 0.001, ** p < 0.01, * p < 0.05,. p < 0.1.

| **Predictor** | **β** | **SE** | **t** | **p** | **R²** |
|---|---|---|---|---|---|
| **A)** Artemia survival | 0.20704 | 0.14603 | 1.418 | 0.1703 | 0.08372 |
| **B)** *Dunaliella* growth rate | 0.44279 | 0.08943 | 4.951 | 5.93e-05 *** | 0.527 |
| **C)** Artemia survival × *Dunaliella* growth rate | 0.79080 | 0.24507 | 3.227 | 0.00388 ** | 0.3212 |

**Supplementary Table 4:**
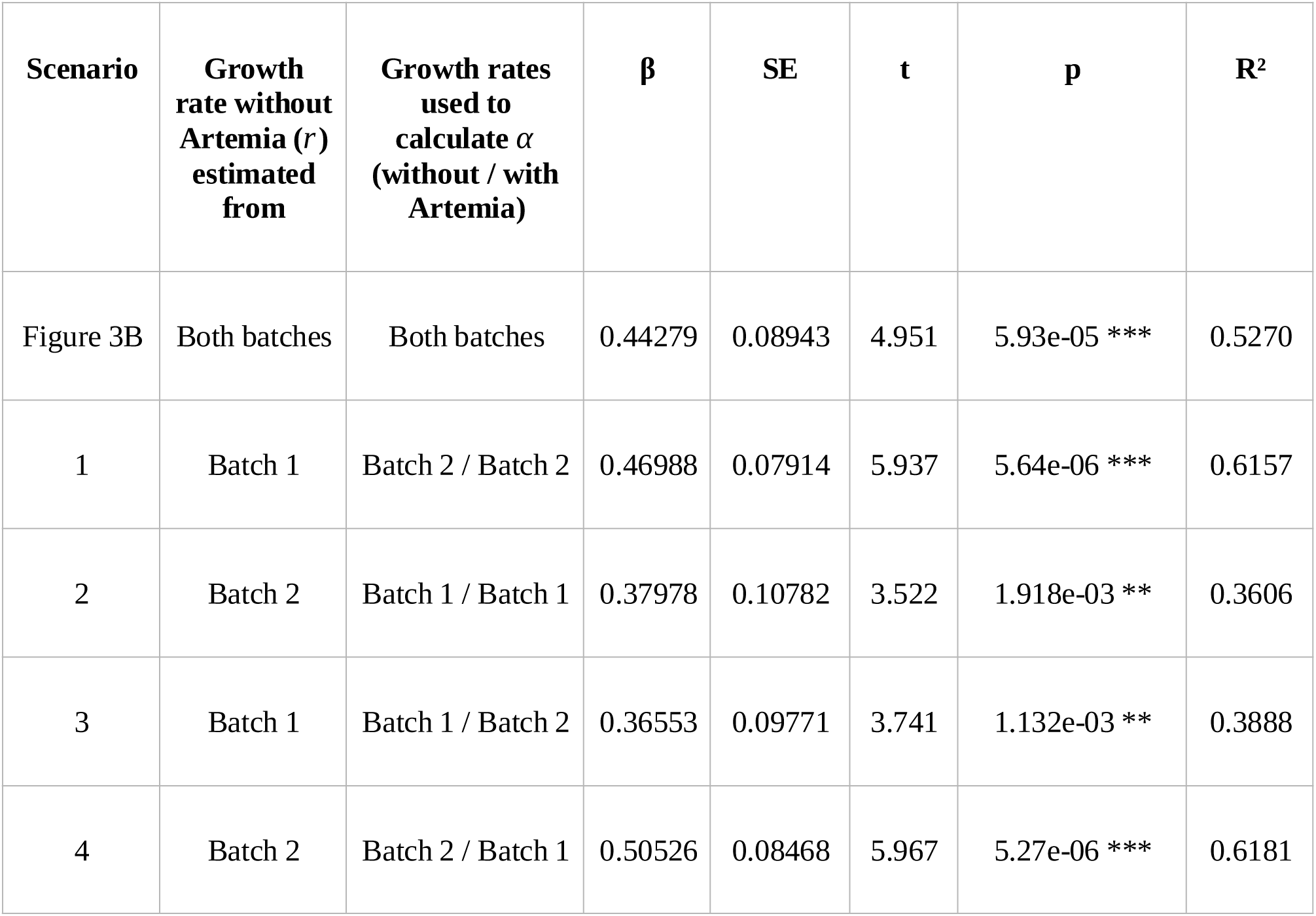
Sensitivity analyses of the relationship between predicted predation rate (*α*) and *Dunaliella* growth rate (*r*). The growth rate without *Artemia* is estimated from one batch, while *α* is calculated as the difference between growth rates without and with Artemia, divided by 5. In scenarios 1 and 2, both growth rates used to calculate *α* are taken from the same batch, whereas in scenarios 3 and 4, the growth rates without and with Artemia are taken from different batches, as indicated in the table. For each model, regression coefficients (β), standard errors (SE), t-values, and associated p-values are reported. Model fit is reported using the coefficient of determination (RZ). Significance levels are indicated as follows: *** p < 0.001, ** p < 0.01, * p < 0.05,. p < 0.1.

## Notes

https://gitlab.com/Louguyot/env_tolerance_niche_realization_guyot_2026

### Competing Interest Statement

The authors have declared no competing interest.

### Summary of Updates

Revised version following the reviewers suggestions and additional revisions. Figures, tables, the title, and the main text have been modified.

